# Peptide-based ligand antagonists block a *Vibrio cholerae* adhesin

**DOI:** 10.1101/2025.09.08.674952

**Authors:** Mingyu Wang, Grace Du, Charity Yongo-Luwawa, Angie Lu, Brett Kinrade, Kim Munro, Karl E. Klose, William D. Lubell, Peter Davies, Shuaiqi Guo

## Abstract

*Vibrio cholerae*, the causative agent of cholera, uses surface proteins such as the repeats-in-toxin (RTX) adhesin FrhA to colonize hosts and initiate infection. Blocking bacterial adhesion represents a promising therapeutic strategy to treat infections without promoting drug resistance. FrhA contains a peptide-binding domain (PBD) that is key for hemagglutination, human epithelial cell binding, and *V. cholerae* biofilm formation. Previous studies identified a lead pentapeptide ligand with the sequence Ala-Gly-Tyr-Thr-Asp (AGYTD) that blocks *V. cholerae* colonization of the mouse small intestine at high micromolar concentrations. A structure-guided approach has now identified a minimal D-amino acid-containing tripeptide motif with higher affinity for the FrhA-PBD and predicted metabolic stability. Our results contribute to the development of anti-adhesion strategies to combat infections.

## Introduction

Bacterial adhesion to host tissues is typically the first step in establishing infections [1]. Without attachment to host tissues, bacteria are rapidly cleared by fluid flow. Stable attachment to the host tissues is typically essential for subsequent colonization and virulence [2-4]. Bacterial adhesion is mediated by surface-exposed adhesion proteins which may assist in the evasion of host immune defenses [3, 5, 6].

The Gram-negative bacterium *Vibrio cholerae* is the causative agent of the diarrheal disease cholera, which is responsible for over 100,000 deaths globally each year [7]. Multiple adhesins in *V. cholerae* contribute to attachment to the intestinal epithelium [8]. For example, the repeats-in-toxin (RTX) adhesin FrhA promotes binding to chitin, erythrocytes, and epithelial cells [5, 9], and assists in biofilm formation [10]. FrhA is localized on the outer membrane of *V. cholerae* by the Type I Secretion System [5, 9, 11]. The retention of FrhA on the surface is regulated by a periplasmic proteolysis system through the signaling molecule c-di-GMP [12-15]. The *N*-terminus of FrhA helps anchor the adhesin to the bacterial surface [16]. The *C*-terminal ligand-binding region (LBR) projects away from the bacterial surface for binding other cells and surfaces (Fig. 1A) [9]. The LBR of FrhA includes a carbohydrate-binding module (CBM) [17] and a peptide-binding domain (PBD) [10]. The recombinant FrhA-CBM demonstrates hemolytic activity against human type O erythrocytes [17]. The FrhA-PBD is required for hemagglutination by *V. cholerae* O395 and enhances biofilm formation [10]. The two ligand-binding domains engage distinct host receptors and may act synergistically to enhance adhesion and virulence [10, 17, 18].

**Figure 1.**
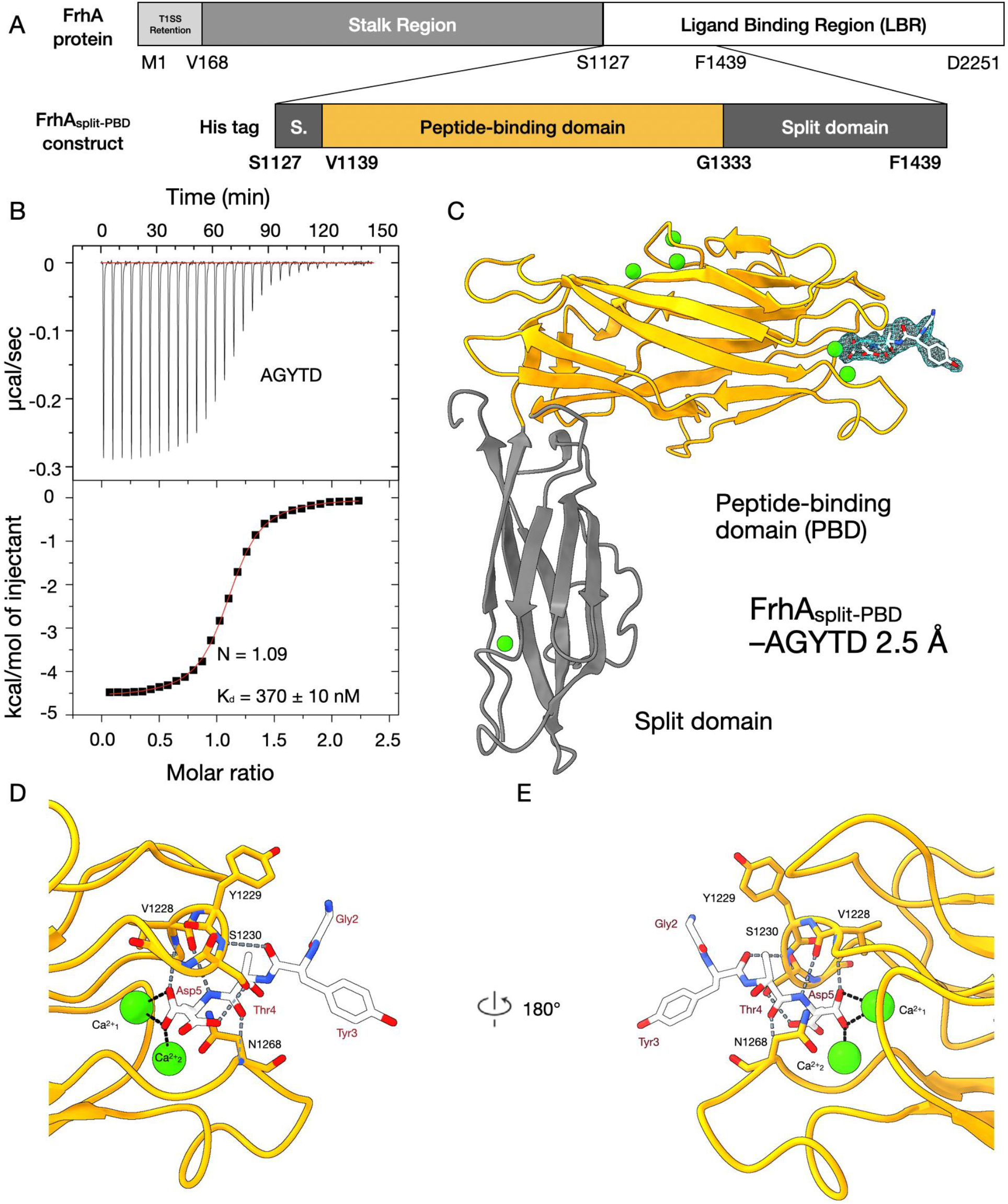
X-ray crystal structure of FrhA_Split-PBD_ revealed interactions with AGYTD. **(A)** The linear domain maps of full-length FrhA and the FrhASplit-PBD construct. The PBD is shown in orange; residue numbers marking domain boundaries are indicated in bold. **(B)** ITC measurement of AGYTD binding to FrhA_Split-PBD_. Fitting of the binding isotherm yielded an N value of 1.09 and a dissociation constant (K_d_) of 370 ± 10 nM. **(C)** The 2.3-Å X-ray crystal structure of FrhA_Split-PBD_ bound to AGYTD (PDB ID: XXXX). The PBD is colored orange, and the companion split domain is colored gray. Ca^2+^ ions are depicted as green spheres, oxygen atoms in red, and nitrogen atoms in blue. The experimental electron density for AGYTD is shown as a light-blue mesh. **(D)** A close-up view of the ligand-binding pocket of FrhA_Split-PBD_, shown in orange, bound to AGYTD. The peptide AGYTD is shown in white, with oxygen atoms in red and nitrogen atoms in blue. Hydrogen bonds are shown as gray dashed lines, and ionic bonds between Ca^2+^ ions (green) and oxygen atoms are shown as black dashed lines. Atomic models of amino acid residue hydrogen-bonds to AGYTD were overlaid on the ribbon model. The *N*-terminal alanine residue of AGYTD was not clearly resolved. Protein residues are labeled with one-letter codes and sequence positions (black), while peptide residues are labeled with three-letter codes (red). **(E)** Back view of the binding pocket shown in (D) following a 180° rotation around the vertical axis.

Anti-adhesion therapies can target pathogen–host attachment by using inhibitors that block specific bacterial adhesins [19, 20]. By preventing host tissue engagements, the inhibitors can impede initial colonization that leads to pathogenesis [19, 21]. Moreover, the inhibitors can also promote the dissociation of bacteria that are already attached to the host tissues and thereby attenuate virulence, because many toxins and effector systems are highly expressed in the adherent state [1, 3, 21-23]. Consequently, adhesin inhibitors facilitate pathogen clearance while reducing local toxin burden [19, 20]. The anti-adhesion strategy has shown efficacy in multiple models against diverse bacterial infections and may impose less selection pressure for antimicrobial resistance than conventional antibiotics [4, 19, 24].

In *V. cholerae*, several lines of evidence support the feasibility of the adhesin inhibitor strategy. Soluble chitooligosaccharides blocked *V. cholerae* from adhering to human colon epithelial cells (HT-29) [25], likely by targeting the *Vibrio* adhesin GbpA [26]. Similarly, oligosaccharides from human milk have been found to inhibit *V. cholerae* adhesion to human erythrocytes and intestinal epithelial cells (Caco-2) [27, 28]. Moreover, L-fucose-based synthetic inhibitors have been shown to mitigate *V. cholerae* adhesion to the villi of the rabbit intestinal epithelium [29]. The inhibitory effects of L-fucose on adhesion *in vivo* are supported and explained by the mechanistic finding that the CBM of FrhA specifically binds to fucosylated glycans [17]. Collectively, carbohydrate adhesin inhibitors have shown potential for reducing adhesion, but do not completely abolish attachment. Targeting additional adhesins and their ligand-binding domains, such as FrhA-PBD, could improve the therapeutic effectiveness of anti-adhesion therapies against *V. cholerae*.

In designing the first inhibitors of FrhA-PBD, we leveraged knowledge of the homologous adhesin *Mp*IBP that binds the Antarctic bacterium *Marinomonas primoryensis* to diatoms and ice [10, 30]. Structure-guided design and iterative screening demonstrated that the peptide-binding domain of *Mp*IBP (*Mp*IBP-PBD), which shares over 60% sequence identity with FrhA-PBD, preferentially binds to the *C*-terminal residues of pentapeptides: Ala-Gly-Tyr-Thr-Asp (AGYTD, dissociation constant K_d_ ≈ 30 nM), Ala-Gly-Tyr-Thr-Ser (AGYTS, K_d_ ≈ 26 nM), and Ala-Gly-Trp-Thr-Asp (AGWTD, K_d_ ≈ 23 nM) [30]. Co-crystal structures of *Mp*IBP-PBD bound to AGYTD revealed that the C-terminal aspartate residue is coordinated by two Ca^2+^ ions in the binding pocket through the terminal α-carboxylate oxygens [30]. The penultimate threonine (the second residue from the *C*-terminus) and the pre-penultimate tyrosine residues (the third residue from the *C*-terminus) of the peptide engage *Mp*IBP-PBD via hydrogen-bond networks. Additionally, the side chain of the peptide pre-penultimate tyrosine interacts with the *Mp*IBP-PBD tyrosine (Y294) via hydrophobic π–π stacking [30, 31]. Considering FrhA-PBD shares high overall sequence identity with *Mp*IBP-PBD with conserved residues lining the binding pocket (Fig. S1), AGYTD was repurposed to target FrhA-PBD [10, 32, 33]. Binding assays *in vitro* confirmed that AGYTD inhibited FrhA-mediated adhesion to human red-blood cells and epithelial (HEp2) cells and disrupted biofilm formation of *V. cholerae* [10]. Furthermore, in the infant mouse competition assay, 500 μM AGYTD showed moderate inhibitory effects on colonization of the small intestine by *V. cholerae* [10].

The relatively low efficacy and poor biological stability of short peptide inhibitors such as AGYTD pose major challenges for developing pharmacological candidates. The high concentration required for effective inhibition (500 µM) would be challenging to maintain in the human intestinal tract, which is enriched in proteases and peptidases capable of rapidly cleaving peptides with natural L-configuration [10, 34]. Solving the structure of FrhA-PBD in complex with its ligands offers a unique opportunity for guiding the rational design of inhibitors with greater binding affinity. Here, we determined the molecular basis of FrhA-PBD-peptide interaction and developed stronger binders. Moreover, based on rational design, amino acids of D-configuration were introduced into the peptide to predictably improve metabolic stability [35]. Our results provide a foundation for developing potent inhibitors to block *V. cholerae* adhesion for attenuating virulence while lowering the risk of developing antimicrobial resistance [4, 19, 24].

## Methods

### Expression and purification of the FrhA_Split-PBD_ construct

The gene encoding the FrhA_Split-PBD_ construct was cloned between the *NdeI* and *XhoI* sites of a pET28a expression vector possessing an *N*-terminal 6× His-tag on the protein. Transformed *Escherichia coli* BL21DE3 (ThermoFisher) cells were grown in LB medium at 37 ºC and subsequently at 23 ºC in the presence of 1mM IPTG (Bioshop) for protein overexpression as previously described [17, 36]. The protein was purified based on previously published protocols [17, 36], using a His GraviTrap™ (Cytiva) Ni^2+^ affinity chromatography column and a Superdex™ 200 Increase 10/300 GL (Cytiva) size-exclusion column.

### Peptide synthesis

All peptides used in X-ray crystallography, ITC measurements, and MST experiments were purchased from GenicBio (Shanghai, China).[30] Representative data from mass spectrometry performed by the vendor support purity greater than 96% (Fig. S2).

### Isothermal titration calorimetry

Isothermal titration calorimetry (ITC) measurements were performed at 20 °C using an iTC_200_ microcalorimeter (Malvern Instruments, Northampton, MA). FrhA_Split-PBD_ was dialyzed overnight against a buffer (50 mM Tris-HCl, pH 9, 150 mM NaCl, 5 mM CaCl_2_) and diluted to a concentration of 20 μM. LyophilizedAGYTD was weighed and solubilized in the same buffer to a concentration of 200 μM.

Each titration consisted of 29 injections. For each injection, 1.3 μL of 200 μM peptide AGYTD solution was added into 20 μM FrhA_Split-PBD,_ using 700 RPM stirring and a 180-second interval between injections. The resulting binding isotherm was fitted using Origin 7.0 software (OriginLab Corp., Northampton, MA) to determine the binding parameters for the interaction.

### Microscale thermophoresis

Microscale thermophoresis (MST) was performed in the presence of the fluorescently labeled FrhA_Split-PBD_ employing unlabeled ligands such as AGYTD. FrhA_Split-PBD_ was labeled using the primary amine-based labeling kit of Red-NHS 2nd Generation (NanoTemper Technologies, San Francisco, CA, USA). The labeling reaction was performed in MST buffer (50 mM HEPES pH 8, 150 mM NaCl, 5 mM CaCl_2_) with a protein concentration of 20 µM (molar ratio of dye:protein ≈ 3:1). The reaction mixture was incubated at room temperature for 1 h in a dark environment for protection from photobleaching. Subsequently, the unreacted dye was removed from the labeled protein using the supplied desalting column, which was equilibrated with MST buffer. The degree of labeling (DOL) was determined using UV/Vis spectrophotometry at 650 and 280 nm—a DOL of 0.5 was typically achieved.

After dialysis of the labeled protein against a buffer containing 50 mM Tris-HCl pH 9, 150 mM NaCl, 5 mM CaCl_2_, and 0.05% v/v Tween-20, the post-dialysis buffer was filtered and used to solubilize the ligands. A series of sixteen 1:1 dilutions of the ligands was prepared, resulting in ligand concentrations ranging from 440 μM to 13.4 nM. After 1 h incubation, the samples were loaded into standard monolith NT.115 capillaries from NanoTemper Technologies. The MST measurements were conducted using a Monolith NT.115 instrument at an ambient temperature of ∼ 22 °C, with instrument parameters set to 20% excitation power and 40% MST power. Three independently prepared replicates were performed for each ligand. The NanoTemper MO.Affinity Analysis software (version 2.3, NanoTemper Technologies) was used to analyze the fluorescence data, fit sigmoidal binding models, and provide the dissociation constants. In the software, the K_d_ values were determined by plotting the change in normalized fluorescence [ΔF_norm_(‰) = F_1_/F_0_] against the logarithm of the concentrations of peptide ligands, where F_1_ and F_0_ respectively correspond to the heated and baseline regions of the thermophoresis traces [37]. The MST on-time, the time between F_1_ and F_0_, was set to 20 seconds for analysis.

### X-ray Crystallography and Data Collection

The FrhA_Split-PBD_ protein construct was respectively incubated with ligand AGYTD and AGWTD in 0.16 M calcium acetate, 0.08 M sodium cacodylate, and 24.4% (w/v) PEG 8000. The protein construct produced thick, triangular crystals after one week. Diffraction data were collected at the CMCF-IM beamline of the Canadian Light Source synchrotron. The phase problem was solved using molecular replacement with the ∼ 60%-identical *Mp*IBP-PBD. The FrhA_Split-PBD_ protein construct corresponds to residues S1127 to F1339 in the full-length FrhA sequence.

### Molecular model building

Molecular replacement of the FrhA_Split-PBD_ was performed using the structure of the *Mp*IBP-PBD (PDB ID: 6X5W). The initial models were improved by rounds of refinement in Phenix [38, 39] and manual model building in Coot [40, 41]. Residues S1127 and F1439 were modeled in the final X-ray crystal structures [10]. Figures relevant to structural studies were prepared using ChimeraX 1.9 [42, 43].

## Results

### The pentapeptide AGYTD binds the PBD of FrhA with nanomolar affinity

Previously, we demonstrated that the pentapeptide AGYTD inhibits *V. cholerae* hemagglutination [30] and intestinal colonization, albeit at relatively high micromolar concentrations [10]. To determine whether AGYTD directly binds the PBD of FrhA, *in vitro* characterization of the FrhA-PBD is required. Because the isolated single-domain FrhA-PBD exhibited a strong tendency to aggregate, we recombinantly produced a 38-kDa two-domain construct (FrhA_Split-PBD_) containing the PBD and the adjacent “split” domain to reduce the propensity for aggregation (Fig. 1A). The *N*-terminally His-tagged FrhA_Split-PBD_ was purified to homogeneity through a combination of Ni^2+^-NTA affinity chromatography and size-exclusion chromatography.

To validate the direct binding of AGYTD to FrhA_Split-PBD_, we performed isothermal titration calorimetry (ITC). The peptide bound FrhA_Split-PBD_ with submicromolar affinity (K_d_ of ∼ 370 ± 10 nM) in a 1:1 stoichiometry (Fig. 1B). The stoichiometry is consistent with the structural data from the homologous *Mp*IBP, which contains a single Ca^2+^-dependent peptide-binding site.

### The structure of FrhA_Split-PBD_-AGYTD complex reveals the molecular basis of protein-peptide interactions

Although AGYTD binds FrhA_Split-PBD_ with submicromolar affinity, high micromolar concentrations of the peptide are required *in vivo* to modestly reduce *V. cholerae* colonization in the mouse intestine [10]. This limitation underscores the need for inhibitors with higher potency as well as greater metabolic stability to withstand the peptidases and proteases in intestinal fluid. To gain structural insights that could help guide such improvements, we crystallized FrhA_Split-PBD_ in complex with AGYTD and analog AGWTD, respectively, using under-oil microbatch methods [30]. We solved the structures of these FrhA_Split-PBD_-peptide complexes using molecular replacement with *Mp*IBP domains as search models. The FrhA_Split-PBD_-AGYTD complex was resolved to 2.4 Å resolution in the space group of P1, with six symmetry-related molecules in the unit cell (Fig. 1C). The split and PBD domains adopted the same overall fold as those in *Mp*IBP, as both formed oblong β-sandwich structures. The cores of the domains were composed mainly of antiparallel β-strands with flexible loops at the extremities. Each protein chain contained six Ca^2+^ ions: two in the ligand-binding pocket, three at another site in the PBD, and one buried in the split domain.

The overall mode of peptide engagement was conserved between the homologous *Mp*IBP-PBD and FrhA-PBD binding pockets. Key interactions included ionic coordination between the two PBD Ca^2+^ ions and the carboxylate group of the C-terminal aspartate (AGYT**<u>D</u>**, Fig. 1C–E) as well as hydrogen bonding from the penultimate threonine hydroxyl group (AGY**<u>T</u>**D) [30]. However, notable differences were observed at the pre-penultimate position: the *Mp*IBP-PBD*–*AGYTD structure showed that the pre-penultimate tyrosine residue (AG**<u>Y</u>**TD) projected upward to form stabilizing π–π interactions with a conserved tyrosine (Y294) near the binding pocket (Fig. S3) [30]. In contrast, in FrhA_Split-PBD_ the peptide tyrosine residue points downward and does not participate in hydrophobic stacking (Fig. 1C, Fig. S4). Furthermore, the glycine residue at the fourth position (A**<u>G</u>**YTD) remains solvent-exposed and did not interact with the protein. The alanine residue at the fifth position (**<u>A</u>**GYTD) was not resolved, suggesting that this residue may be flexible or disordered (Fig. 1C). Together, these observations indicated that the two *N*-terminal residues of AGYTD may be dispensable for binding, and that shorter peptide fragments may retain affinity while reducing molecular complexity.

### Tripeptide YTD is a stronger binder than the pentapeptide AGYTD

Considering that structural analysis indicated that the binding pocket engaged only the three C-terminal residues of AGYTD, we tested shorter peptide variants (GYTD, YTD, and TD; GenicBio) for affinity to FrhA_Split-PBD_. To circumvent the requirement for large quantities of pure protein and ligands for the labor-intensive ITC setup [37], we used micro-scale thermophoresis (MST) as a higher-throughput alternative. MST determines binding by detecting fluorescence changes in a temperature gradient and requires only one binding partner to be fluorescently labeled. In our assay, FrhA_Split-PBD_ was fluorescently labeled mostly on the primary amines of lysine residues located away from the PBD binding site using NHS-ester chemistry. To validate this workflow, we first measured binding of AGYTD, which yielded a K_d_ of ∼ 500 ± 140 nM, consistent with the ITC results (370 ± 10 nM). Among the truncated variants, the tetrapeptide GYTD bound weaker (K_d_ ≈ 1.1 ± 0.37 µM), but the tripeptide YTD bound significantly more tightly (K_d_ ≈ 190 ± 62 nM) than AGYTD (Fig. 2A–B, Fig. S5). In contrast, truncation to the dipeptide TD abolished binding altogether. These results highlight the nuanced effects of N-terminal truncation on AGYTD and demonstrate that the removal of specific residues can either weaken or enhance affinity. Notably, the stronger binding of YTD compared to AGYTD identifies a minimal tripeptide motif that maintains high affinity while reducing molecular complexity, providing a promising template for small-molecule mimics and synthetic optimization.

**Figure 2.**
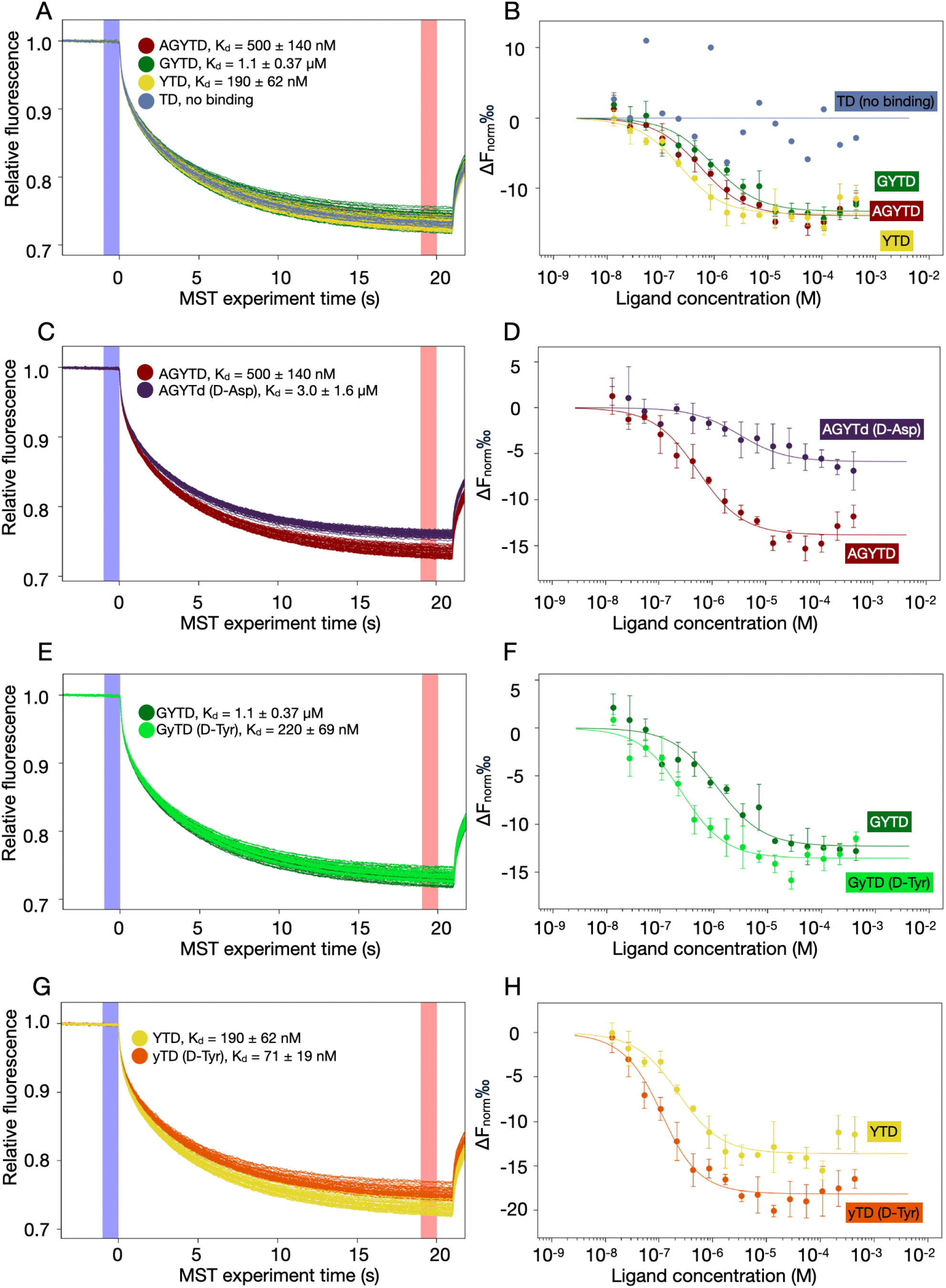
Binding of FrhA_Split-PBD_ to the pentapeptide AGYTD and variants measured by MST. **(A–H)** Thermophoretic mobility traces and dose-response binding curves for reactions of FrhA_Split-PBD_ binding to various groups of ligands: AGYTD (dark red), GYTD (dark green), YTD (yellow), TD (blue), AGYTd (purple), GyTD (light green), and yTD (orange). **(A, C, E, and G)** Thermophoretic mobility traces of the MST reactions depict individual traces for each ligand concentration and highlight the F_0_ (blue) and F_1_ (red) regions used to calculate binding. **(B, D, F, and H)** Dose–response binding curves for fluorescently labeled FrhA_Split-PBD_ with the ligands. Data points are the mean from three independent experiments, plotted as ΔF_norm_ (F_1_/F_0_) versus ligand concentration (in M). Error bars show standard deviation. Solid lines were fitted using the law of mass action to derive K_d_ values.

### Replacement of L-tyrosine with its D-enantiomer at the pre-penultimate position enhances ligand affinity for the PBD

Having established that the tripeptide YTD improved binding affinity, we sought to enhance peptide stability against enzymatic degradation in the gastrointestinal tract. To resist hydrolysis by carboxypeptidases [44], which cleave peptides from the *C* termini, we substituted the terminal L-aspartic acid with its D enantiomer. This modification yielded the peptide Ala-Gly-Tyr-Thr-D-Asp (AGYT**<u>d</u>**; lowercase letter indicates residues in D configuration), which bound FrhA_Split-PBD_ with significantly reduced affinity compared to AGYTD (K_d_ ≈ 3.0 ± 1.6 µM vs 500 ± 140 nM; Fig. 2C–D). This loss of affinity may have been expected, because the stereochemical inversion reorientated the terminal aspartate side chain and likely disrupted coordinating hydrogen bonds in the binding pocket.

We then tested whether stereochemical modification at the pre-penultimate residue could enhance binding. Substituting L- with D-tyrosine in the tetrapeptide GyTD was hypothesized to reorient the aromatic side chain, enabling stronger π–π stacking with Y1229 in FrhA_Split-PBD_ (Fig. 2E–F). Indeed, this substitution improved binding by approximately sevenfold with a K_d_ of ∼ 220 ± 69 nM from GYTD (K_d_ ≈ 1.1 ± 0.37 µM).

Encouraged by this result, we next examined the tripeptide yTD by removing the N-terminal glycine residue (Fig. 2G–H). MST measurements showed that yTD binds with a K_d_ of ∼ 71 ± 19 nM, greater than two-fold stronger than that of YTD (K_d_ ≈ 190 ± 62 nM) and greater than seven-fold stronger than the initial pentapeptide lead AGYTD. This single modification conferred two key advantages: enhanced inhibitory potency and predicted resistance to degradation by aminopeptidases and related enzymes due to the D-amino acid at the *N*-terminus [45].

Having established the affinity-enhancing effects of truncation to a tripeptide and stereochemical inversion of the aromatic amino acid residue, we next asked whether similar modifications could enhance binding in a different ligand. To explore this, we turned to AGWTD (K_d_ ≈ 380 ± 100 nM to FrhA_Split-PBD_), another high-affinity pentapeptide binder of *Mp*IBP-PBD, for truncation and D-amino acid substitution. Structural analysis of the FrhA_Split-PBD_-AGWTD complex revealed a peptide conformation like that of AGYTD (Fig. 3A–B). Notably, no π-π stacking was observed between the peptide tryptophan and Y1229 of the protein. Truncating AGWTD to the tripeptide WTD increased binding affinity to a K_d_ of ∼ 200 ± 46 nM compared to the parent pentapeptide (K_d_ ≈ 380 ± 100 nM, Fig. 3C–D). Substituting the tryptophan in the tripeptide with its D-enantiomer further strengthened binding, improving the affinity to a K_d_ of ∼ 120 ± 41 nM.

**Figure 3.**
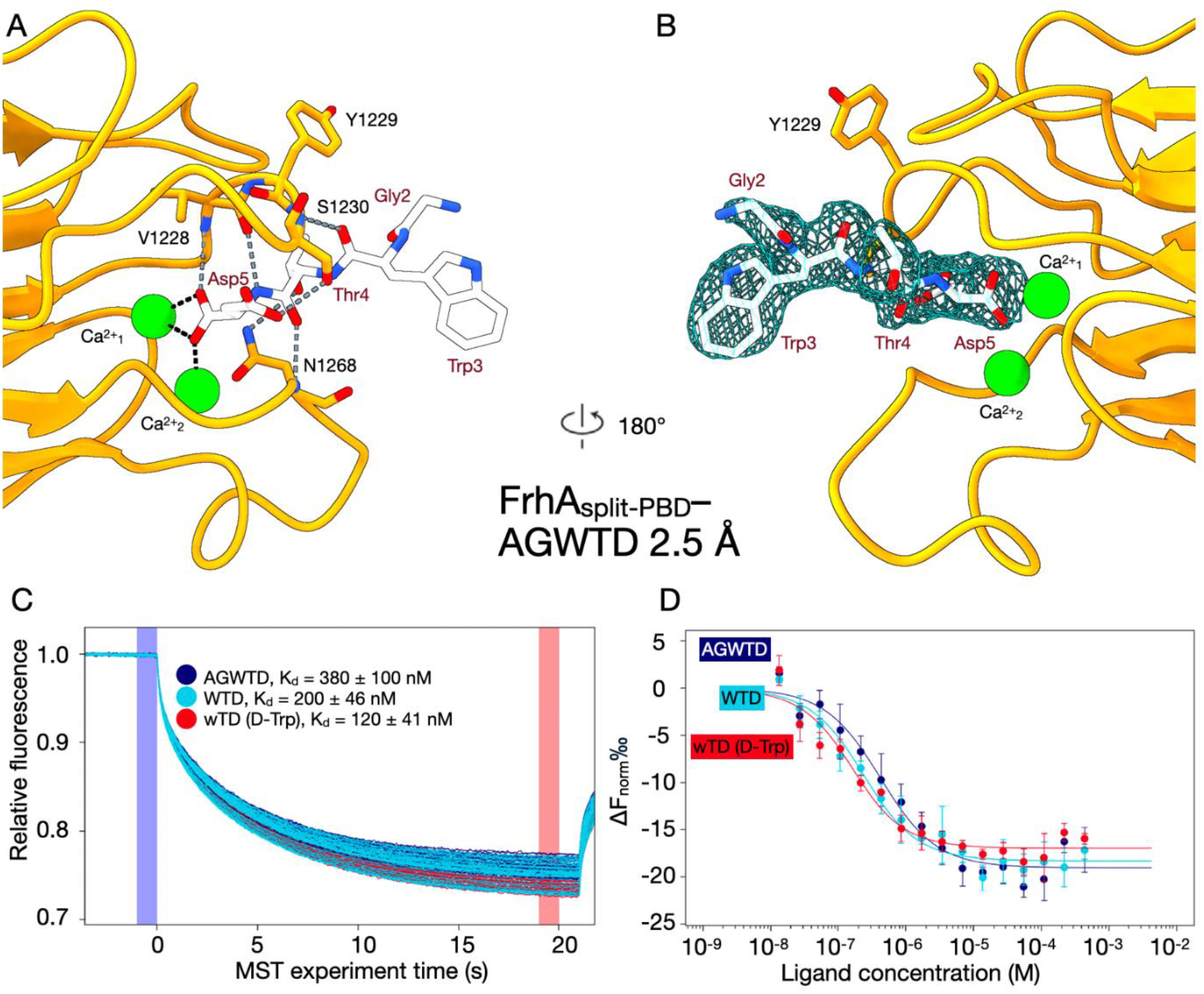
Binding of FrhA_Split-PBD_ to the pentapeptide AGWTD and variants measured by MST. **(A)** Zoomed-in views of the ligand binding site of the 2.5-Å X-ray crystal structure of the FrhA_Split-PBD_**–**AGWTD complex (PDB ID: XXXX). The PBD is shown in orange, Ca^2+^ ions in green, and the peptide in white. The same color scheme was used as shown in in Fig.1. **(B)** Back view of the binding pocket shown in (A) after a 180° rotation around the vertical axis. The experimental electron density of AGWTD is shown as a light-blue mesh. **(C–D)** Thermophoretic mobility traces and dose-response binding curves for reactions of FrhASplit-PBD binding AGWTD (dark blue), WTD (light blue), and wTD (red). **(C)** Thermophoretic mobility traces of the MST reactions depict individual traces for each ligand concentration and highlight the F_0_ (blue) and F_1_ (red) regions used to calculate binding. **(D)** Dose–response binding curves for fluorescently labeled FrhA_Split-PBD_ with representative ligands. Data points are the mean from three independent experiments, plotted as ΔF_norm_ (F_1_/F_0_) versus ligand concentration (in M). Error bars show standard deviation. Solid lines were fitted using the law of mass action to derive K_d_ values.

**Figure 4.**
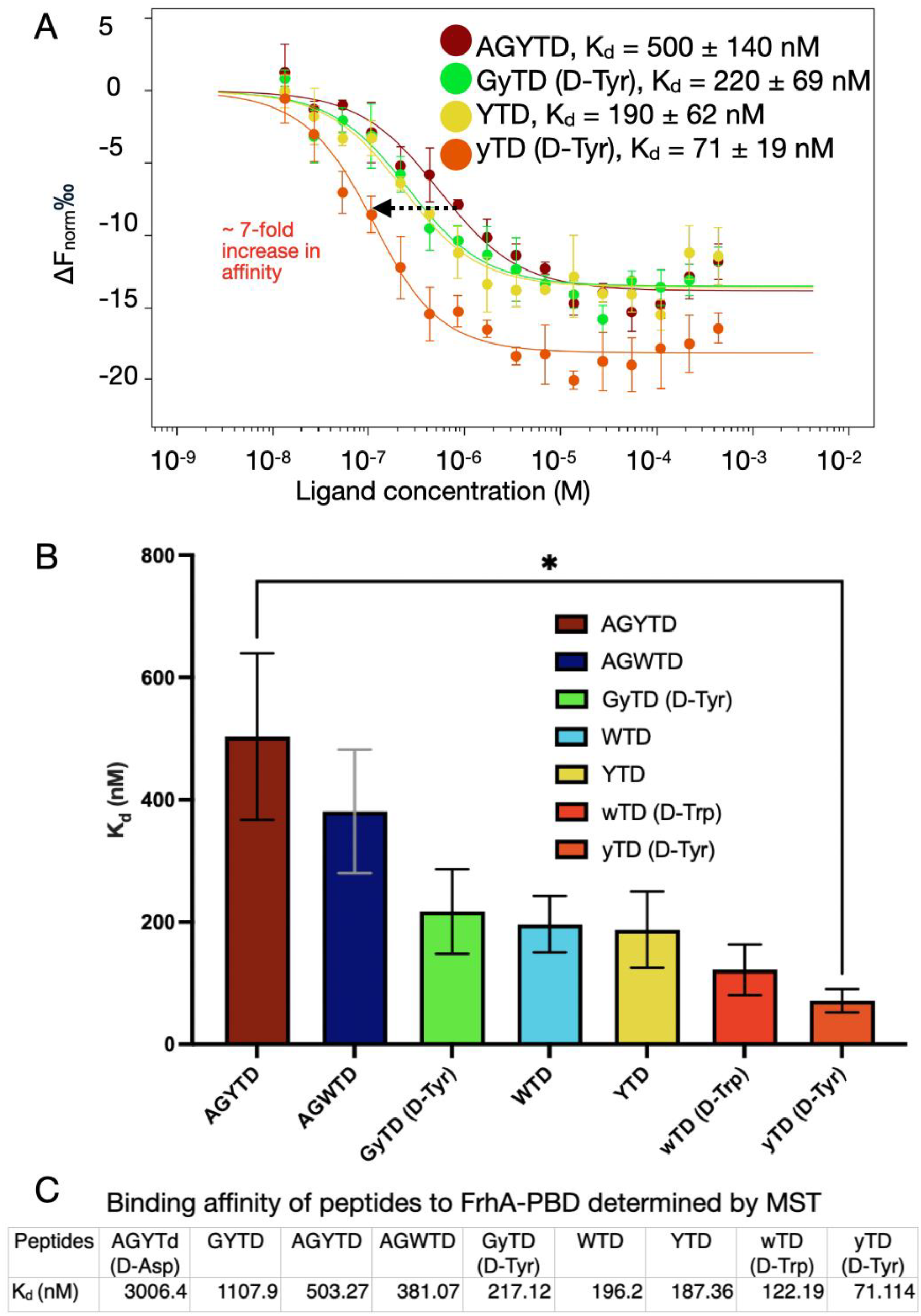
Tripeptide yTD showed a sevenfold increase in affinity compared to the parent pentapeptide AGYTD. **(A)** Dose–response binding curves for fluorescently labeled FrhA_Split-PBD_ with the ligands AGYTD (dark red), GyTD (light green), YTD (yellow), and yTD (orange), as shown in previous figures. Data points are the mean from three independent experiments, plotted as ΔF_norm_ (F_1_/F_0_) versus ligand concentration (in M). Error bars show standard deviation. Solid lines were fitted using the law of mass action to derive K_d_ values. **(B)** Summary of the binding affinities of peptidyl ligands to FrhA_Split-PBD_. The peptides were shown with decreasing numbers of amino acid residues from left to right. Error bars show K_d_ confidence interval (1 standard deviation). The bracket with an asterisk indicates the statistically significant difference in the binding affinities of AGYTD and yTD determined by an unpaired t-test with a false discovery rate set below 0.01. **(C)** Table displaying the binding affinity of peptides to FrhA-PBD and the 68% confidence interval determined by three independent MST experiments each.

Together, these results establish that D-amino acid substitution at the pre-penultimate position can enhance both affinity and potential proteolytic stability across multiple peptide series. The identification of compact tripeptide motifs such as D-Tyr-Thr-Asp and D-Trp-Thr-Asp highlights a general strategy for creating small scaffolds that bind the FrhA_Split-PBD_ with high-affinity and lays a foundation for developing peptidomimetics and non-peptide inhibitors of the PBD with improved pharmacokinetic properties.

## Discussion

Despite decades of progress, the development of anti-adhesion treatments remains challenging [20, 21]. To have clinical utility, an inhibitor must target the adhesin with high specificity and binding affinity, as well as remain active at physiologically relevant concentrations by resisting degradation in the protease-rich intestinal environment [19, 34]. In this study, we systematically addressed these key challenges by optimizing peptide inhibitors for FrhA-PBD. Using a recombinant FrhA_Split-PBD_ construct, we validated direct peptide binding *in vitro*, complementing previous *in vivo* evidence from the infant mouse model [10]. X-ray crystal structures of FrhA_Split-PBD_ in complex with ligands guided the rational design of shorter derivatives with improved affinity. In parallel, the selective incorporation of D-amino acids further enhanced potency and provided a strategy to improve predicted inhibitor stability against proteases [35].

To establish a quantitative framework for comparing peptide performance, we first measured the affinity of the starting peptide AGYTD by ITC. For subsequent inhibitor optimization, we prioritized MST because it requires far less protein than ITC and enabled rapid and reproducible results for screening the relative potency of peptide variants [37]. Systematic truncation of AGYTD into GYTD, YTD, and TD revealed a non-linear relationship between peptide length and binding affinity; GYTD exhibited weaker binding than AGYTD (Fig. 2A–B).

Tripeptide YTD exhibited strong binding to FrhA_Split-PBD_. Removal of the pre-penultimate tyrosine residue abolished binding in the dipeptide TD (Fig. 2A–B). This loss can be explained by the absence of π-stacking interactions provided by the tyrosine side chain and the elimination of hydrogen bonds from its main-chain carbonyl oxygen [31]. More importantly, TD is substantially more polar than YTD and better solvated by water molecules in solution [46, 47]. For binding to PBD, structured water molecules must be displaced from both the dipeptide TD and the binding pocket of the PBD, a process that incurs a significant enthalpic cost [46, 48]. Consequently, the high desolvation penalty likely makes binding thermodynamically less favorable, explaining the complete loss of interaction [46, 48].

Having established the contributions of peptide length for binding through truncation, we next examined how stereochemical changes at specific positions impact peptide–PBD interactions. The effects of D-amino acid substitution on peptide binding affinity were position-dependent, reflecting the local structural environment within the binding pocket. Substitution of L- for D-aspartic acid at the *C*-terminus greatly reduced binding affinity. This outcome may have been anticipated, because previous fluorescence polarization experiments with the homologous protein *Mp*IBP-PBD showed that certain modifications to the *C*-terminal carboxylate group abolished binding [30]. Our structural data support this interpretation. In the X-ray crystal structure of the FrhA_Split-PBD_–AGYTD complex, the *C*-terminal carboxylate group of the aspartic acid coordinates two Ca^2+^ ions in the binding pocket; one carboxylate oxygen and the backbone carbonyl oxygen of the aspartic acid form hydrogen bonds with the protein residues V1228 and S1230, respectively (Fig. 2D–E). Inverting stereochemistry at this critical Asp residue alters the orientation of both the terminal and side-chain carboxylates, disrupting the geometry required for the bonding network.

In contrast, D-amino acid substitution at the pre-penultimate position (**<u>y</u>**TD) was well tolerated. The tyrosine residue lies further from the core of the binding pocket, is largely solvent-exposed, and contributes only a single hydrogen bond to S1230. As a result, inverting the stereochemistry of this residue minimally affected Ca^2+^coordination as well as the hydrogen-bond network. Instead, enhanced binding affinity was achieved, possibly through improved engagement of Y1229 in the PBD by the D-tyrosine residue.

Future work is merited to optimize the minimal yTD scaffold as a structural template that captures the essential interactions for strong PBD binding. Incorporation of D-amino acids may offer a promising route to enhance proteolytic stability, because antimicrobial peptides possessing D-residues are known to resist cleavage by intestinal proteases such as trypsin [49]. Consistent with other peptides with *N*-terminal D-residues [34, 45], yTD is expected to be resistant to aminopeptidases. In the presence of endopeptidases like α-chymotrypsin, which can digest peptides with both L- and D-Trp, the latter are cleaved more slowly [50]. These precedents suggest that yTD is likely to persist longer in the intestinal lumen [34]. Future studies could experimentally test this prediction using *in vitro* enzyme digestion assays in simulated intestinal fluids coupled with mass spectrometric analysis to quantify degradation and confirm stability.

Looking ahead, future therapeutic strategies may employ a cocktail of adhesin-specific inhibitors to achieve broad-spectrum anti-adhesion coverage across *V. cholerae* biotypes [20, 21]. The epidemiologically relevant El Tor biotype, responsible for the seventh and ongoing cholera pandemic, potentially expresses different adhesins than the classical biotype [51]. The role of FrhA in adhesion in the El Tor biotype remains poorly defined, whereas other adhesins, such as the mannose-sensitive hemagglutinin (MSHA) pilus [51, 52] and the toxin-coregulated pilus (TCP) [53-55], are more prominent in this strain and contribute to virulence. In this context, the inhibitors of FrhA-PBD may serve as one component of a broader anti-adhesion strategy that also includes inhibitors of FrhA-CBM and other adhesins like MSHA, to achieve broad-spectrum anti-adhesion coverage across *V. cholerae* biotypes [20]. Overall, this multi-inhibitor strategy could offer a viable adjunct or alternative to treatment by conventional antibiotics in clinically relevant settings.

## Author contributions

SG, WDL, PLD and MW conceived the research; MW, BK, GD, KM, AL, CY, and SG performed the experiments; MW and SG wrote the manuscript. All authors contributed to the editing and revision of the manuscript.

## Acknowledgements

We thank Dr. Joaquin Ortega and Dominic Arpin for their assistance in training for MST experiments and interpretation of the results. PLD acknowledges funding from CIHR Foundation Award FRN 148422. SG is supported by an NSERC Discovery Grant (RGPIN-2024-04631), a Fonds de recherche du Québec – Santé (FRQS) Research Scholar (Junior 1) award (359456 & 376506), a McGill Centre de Recherche en Biologie Structurale (CRBS) Bluesky award, and a start-up fund by the McGill Faculty of Medicine and Health Sciences. MW is supported by studentships awarded by the CRBS and Canadian Antimicrobial Resistance Network (CAN-AMR-NET). GD was supported by an NSERC Undergraduate Student Research Award (USRA).

## PBD deposition

X-ray crystal structures of FrhA_split-PBD_ in complex with AGYTD and AGWTD were deposited into the Protein Data Bank with access code XXXX and YYYY.

## Data availability statement

All the data provided in the manuscript are available from the corresponding author upon reasonable request.

## References

1. Berne, C., Ducret, A., Hardy, G. G. & Brun, Y. V. (2015) Adhesins involved in attachment to abiotic surfaces by Gram-negative bacteria, Microbial biofilms, 163–199.

2. Patel, S., Mathivanan, N. & Goyal, A. (2017) Bacterial adhesins, the pathogenic weapons to trick host defense arsenal, Biomedicine & Pharmacotherapy. 93, 763–771.

3. Stones, D. H. & Krachler, A. M. (2016) Against the tide: the role of bacterial adhesion in host colonization, Biochemical Society Transactions. 44, 1571–1580.

4. Rasko, D. A. & Sperandio, V. (2010) Anti-virulence strategies to combat bacteria-mediated disease, Nature reviews Drug discovery. 9, 117–128.

5. Syed, K. A., Beyhan, S., Correa, N., Queen, J., Liu, J., Peng, F., Satchell, K. J. F., Yildiz, F. & Klose, K. E. (2009) The Vibrio cholerae Flagellar Regulatory Hierarchy Controls Expression of Virulence Factors, Journal of Bacteriology. 191, 6555–6570.

6. Hancock, V., Witsø, I.L. & Klemm, P. (2011) Biofilm formation as a function of adhesin, growth medium, substratum and strain type, International Journal of Medical Microbiology. 301, 570–576.

7. Ali, M., Lopez, A. L., You, Y. A., Kim, Y. E., Sah, B., Maskery, B. & Clemens, J. (2012) The global burden of cholera, Bulletin of the World Health Organization. 90, 209–218.

8. Almagro-Moreno, S., Pruss, K. & Taylor, R. K. (2015) Intestinal Colonization Dynamics of Vibrio cholerae, PLOS Pathogens. 11, e1004787.

9. Satchell, K. J. (2011) Structure and function of MARTX toxins and other large repetitive RTX proteins, Annual review of microbiology. 65, 71–90.

10. Lloyd, C. J., Guo, S., Kinrade, B., Zahiri, H., Eves, R., Ali, S. K., Yildiz, F., Voets, I. K., Davies, P. L. & Klose, K. E. (2023) A peptide-binding domain shared with an Antarctic bacterium facilitates <i>Vibrio cholerae</i> human cell binding and intestinal colonization, Proceedings of the National Academy of Sciences. 120.

11. Spitz, O., Erenburg Isabelle, N., Beer, T., Kanonenberg, K., Holland, I. B. & Schmitt, L. (2019) Type I Secretion Systems—One Mechanism for All?, Microbiology Spectrum. 7, 10.1128/microbiolspec.psib-0003-2018.

12. Newell, P. D., Boyd, C. D., Sondermann, H. & O’Toole, G. A. (2011) A c-di-GMP Effector System Controls Cell Adhesion by Inside-Out Signaling and Surface Protein Cleavage, PLOS Biology. 9, e1000587.

13. Boyd, C. D., Chatterjee, D., Sondermann, H. & O’Toole, G. A. (2012) LapG, required for modulating biofilm formation by Pseudomonas fluorescens Pf0-1, is a calcium-dependent protease, Journal of Bacteriology. 194, 4406–4414.

14. Newell, P. D., Monds, R. D. & O’Toole, G. A. (2009) LapD is a bis-(3′,5′)-cyclic dimeric GMP-binding protein that regulates surface attachment by <i>Pseudomonas fluorescens</i> Pf0–1, Proceedings of the National Academy of Sciences. 106, 3461–3466.

15. Kitts, G., Giglio Krista, M., Zamorano-Sánchez, D., Park Jin, H., Townsley, L., Cooley Richard, B., Wucher Benjamin, R., Klose Karl, E., Nadell Carey, D., Yildiz Fitnat, H. & Sondermann, H. (2019) A Conserved Regulatory Circuit Controls Large Adhesins in Vibrio cholerae, mBio. 10, 10.1128/mbio.02822-19.

16. Smith, T. J., Font Maria, E., Kelly Carolyn, M., Sondermann, H. & O’Toole George, A. (2018) An N-Terminal Retention Module Anchors the Giant Adhesin LapA of Pseudomonas fluorescens at the Cell Surface: a Novel Subfamily of Type I Secretion Systems, Journal of Bacteriology. 200, 10.1128/jb.00734-17.

17. Sherik, M., Eves, R., Guo, S., Lloyd, C. J., Klose, K. E. & Davies, P. L. (2024-02-14) Sugar-binding and split domain combinations in repeats-in-toxin adhesins from Vibrio cholerae and Aeromonas veronii mediate cell-surface recognition and hemolytic activities, mBio. 15.

18. Ye, Q., Eves, R., Vance, T. D. R., Hansen, T., Sage, A. P., Petkovic, A., Bradley, B., Escobedo, C., Graham, L. A., Allingham, J. S. & Davies, P. L. (2025) <i>Aeromonas hydrophila</i> RTX adhesin has three ligand-binding domains that give the bacterium the potential to adhere to and aggregate a wide variety of cell types, mBio. 16, e03158–24.

19. Krachler, A. M. & Orth, K. (2013) Targeting the bacteria–host interface: strategies in anti-adhesion therapy, Virulence. 4, 284–294.

20. Ofek, I., Hasty, D. L. & Sharon, N. (2003) Anti-adhesion therapy of bacterial diseases: prospects and problems, FEMS Immunology & Medical Microbiology. 38, 181–191.

21. Asadi, A., Razavi, S., Talebi, M. & Gholami, M. (2019) A review on anti-adhesion therapies of bacterial diseases, Infection. 47, 13–23.

22. Gardel, C. L. & Mekalanos, J. J. (1996) Alterations in Vibrio cholerae motility phenotypes correlate with changes in virulence factor expression, Infection and Immunity. 64, 2246–2255.

23. Mekalanos, J. J. (1992) Environmental signals controlling expression of virulence determinants in bacteria, Journal of bacteriology. 174, 1–7.

24. Pecoraro, C., Carbone, D., Parrino, B., Cascioferro, S. & Diana, P. (2023) Recent developments in the inhibition of bacterial adhesion as promising anti-virulence strategy, International Journal of Molecular Sciences. 24, 4872.

25. Wang, S., Wang, J., Mou, H., Luo, B. & Jiang, X. (2015) Inhibition of adhesion of intestinal pathogens (Escherichia coli, Vibrio cholerae, Campylobacter jejuni, and Salmonella Typhimurium) by common oligosaccharides, Foodborne pathogens and disease. 12, 360–365.

26. Wong, E., Vaaje-Kolstad, G., Ghosh, A., Hurtado-Guerrero, R., Konarev, P. V., Ibrahim, A. F., Svergun, D. I., Eijsink, V. G., Chatterjee, N. S. & van Aalten, D. M. (2012) The Vibrio cholerae colonization factor GbpA possesses a modular structure that governs binding to different host surfaces, PLoS pathogens. 8, e1002373.

27. Holmgren, J., Svennerholm, A. M. & Ahrén, C. (1981) Nonimmunoglobulin fraction of human milk inhibits bacterial adhesion (hemagglutination) and enterotoxin binding of Escherichia coli and Vibrio cholerae, Infection and Immunity. 33, 136–141.

28. Coppa, G. V., Zampini, L., Galeazzi, T., Facinelli, B., Ferrante, L., Capretti, R. & Orazio, G. (2006) Human Milk Oligosaccharides Inhibit the Adhesion to Caco-2 Cells of Diarrheal Pathogens: Escherichia coli, Vibrio cholerae, and Salmonella fyris, Pediatric Research. 59, 377–382.

29. Jones, G. W. & Freter, R. (1976) Adhesive properties of Vibrio cholerae: nature of the interaction with isolated rabbit brush border membranes and human erythrocytes, Infection and immunity. 14, 240–245.

30. Guo, S., Zahiri, H., Stevens, C., Spaanderman, D. C., Milroy, L.-G., Ottmann, C., Brunsveld, L., Voets, I. K. & Davies, P. L. (2021) Molecular basis for inhibition of adhesin-mediated bacterialhost interactions through a peptide-binding domain, Cell Reports. 37, 110002.

31. Calinsky, R. & Levy, Y. (2024) Aromatic Residues in Proteins: Re-Evaluating the Geometry and Energetics of π–π, Cation−π, and CH−π Interactions, The Journal of Physical Chemistry B. 128, 8687–8700.

32. Madeira, F., Madhusoodanan, N., Lee, J., Eusebi, A., Niewielska, A., Tivey, A. R., Lopez, R. & Butcher, S. (2024) The EMBL-EBI Job Dispatcher sequence analysis tools framework in 2024, Nucleic acids research. 52, W521–W525.

33. Sievers, F., Wilm, A., Dineen, D., Gibson, T. J., Karplus, K., Li, W., Lopez, R., McWilliam, H., Remmert, M., Söding, J., Thompson, J. D. & Higgins, D. G. (2011) Fast, scalable generation of high-quality protein multiple sequence alignments using Clustal Omega, Molecular Systems Biology. 7, 539.

34. Accardo, F., Prandi, B., Dellafiora, L., Tedeschi, T. & Sforza, S. (2024) How D-amino acids embedded in the protein sequence modify its digestibility: Behaviour of digestive enzymes tested on a model peptide used as target, Food Chemistry. 458, 140175.

35. Lu, J., Xu, H., Xia, J., Ma, J., Xu, J., Li, Y. & Feng, J. (2020) D-and unnatural amino acid substituted antimicrobial peptides with improved proteolytic resistance and their proteolytic degradation characteristics, Frontiers in Microbiology. 11, 563030.

36. Garnham, C. P., Gilbert, J. A., Hartman, C. P., Campbell, R. L., Laybourn-Parry, J. & Davies, P. L. (2008) A Ca2+-dependent bacterial antifreeze protein domain has a novel β-helical ice-binding fold, Biochemical Journal. 411, 171–180.

37. Gontier, A., Varela, P. F., Nemoz, C., Ropars, V., Aumont-Nicaise, M., Desmadril, M. & Charbonnier, J.-B. (2021) Measurements of Protein–DNA Complexes Interactions by Isothermal Titration Calorimetry (ITC) and Microscale Thermophoresis (MST) in Multiprotein Complexes: Methods and Protocols (Poterszman, A., ed) pp. 125-143, Springer US, New York, NY.

38. Liebschner, D., Afonine, P. V., Baker, M. L., Bunkóczi, G., Chen, V. B., Croll, T. I., Hintze, B., Hung, L.-W., Jain, S. & McCoy, A. J. (2019) Macromolecular structure determination using X-rays, neutrons and electrons: recent developments in Phenix, Biological Crystallography. 75, 861–877.

39. Afonine, P. V., Grosse-Kunstleve, R. W., Echols, N., Headd, J. J., Moriarty, N. W., Mustyakimov, M., Terwilliger, T. C., Urzhumtsev, A., Zwart, P. H. & Adams, P. D. (2012) Towards automated crystallographic structure refinement with phenix. refine, Biological crystallography. 68, 352–367.

40. Emsley, P. & Cowtan, K. (2004) Coot: model-building tools for molecular graphics, Biological crystallography. 60, 2126–2132.

41. Emsley, P., Lohkamp, B., Scott, W. G. & Cowtan, K. (2010) Features and development of Coot, Biological crystallography. 66, 486–501.

42. Goddard, T. D., Huang, C. C., Meng, E. C., Pettersen, E. F., Couch, G. S., Morris, J. H. & Ferrin, T. E. (2018) UCSF ChimeraX: Meeting modern challenges in visualization and analysis, Protein science. 27, 14–25.

43. Pettersen, E. F., Goddard, T. D., Huang, C. C., Meng, E. C., Couch, G. S., Croll, T. I., Morris, J. H. & Ferrin, T. E. (2021) UCSF ChimeraX: Structure visualization for researchers, educators, and developers, Protein science. 30, 70–82.

44. Sung, Y. S., Khvalbota, L., Dhaubhadel, U., Špánik, I. & Armstrong, D. W. (2023) Teicoplanin aglycone media and carboxypeptidase Y: Tools for finding low-abundance D-amino acids and epimeric peptides, Chirality. 35, 461–468.

45. Pearson, R. K., Powers, S. P., Hadac, E. M., Gaisano, H. & Miller, L. J. (1987) Establishment of a new short, protease-resistant, affinity labeling reagent for the cholecystokinin receptor, Biochemical and Biophysical Research Communications. 147, 346–353.

46. Mondal, J., Friesner, R. A. & Berne, B. J. (2014) Role of Desolvation in Thermodynamics and Kinetics of Ligand Binding to a Kinase, Journal of Chemical Theory and Computation. 10, 5696–5705.

47. Olsson, T. S. G., Williams, M. A., Pitt, W. R. & Ladbury, J. E. (2008) The Thermodynamics of Protein–Ligand Interaction and Solvation: Insights for Ligand Design, Journal of Molecular Biology. 384, 1002–1017.

48. Zsidó, B.Z. & Hetényi, C. (2021) The role of water in ligand binding, Current Opinion in Structural Biology. 67, 1–8.

49. Zhao, X., Zhang, M., Muhammad, I., Cui, Q., Zhang, H., Jia, Y., Xu, Q., Kong, L. & Ma, H. (2021) An Antibacterial Peptide with High Resistance to Trypsin Obtained by Substituting d-Amino Acids for Trypsin Cleavage Sites, Antibiotics. 10, 1465.

50. Ingles, D. & Knowles, J. (1967) Specificity and stereospecificity of α-chymotrypsin, Biochemical Journal. 104, 369–377.

51. Silva, A. J. & Benitez, J. A. (2016) Vibrio cholerae Biofilms and Cholera Pathogenesis, PLOS Neglected Tropical Diseases. 10, e0004330.

52. Jones, C. J., Utada, A., Davis, K. R., Thongsomboon, W., Zamorano Sanchez, D., Banakar, V., Cegelski, L., Wong, G. C. & Yildiz, F. H. (2015) C-di-GMP regulates motile to sessile transition by modulating MshA pili biogenesis and near-surface motility behavior in Vibrio cholerae, PLoS pathogens. 11, e1005068.

53. Oki, H., Kawahara, K., Iimori, M., Imoto, Y., Nishiumi, H., Maruno, T., Uchiyama, S., Muroga, Y., Yoshida, A., Yoshida, T., Ohkubo, T., Matsuda, S., Iida, T. & Nakamura, S. (2022) Structural basis for the toxin-coregulated pilus–dependent secretion of <i>Vibrio cholerae</i> colonization factor, Science Advances. 8, eabo3013.

54. Nguyen, M., Wu, T.-H., Danielson Katie J., Khan Nabeel M., Zhang John Z. & Craig, L. (2023) Mechanism of secretion of TcpF by the <i>Vibrio cholerae</i> toxin-coregulated pilus, Proceedings of the National Academy of Sciences. 120, e2212664120.

55. Taylor, R. K., Miller, V. L., Furlong, D. B. & Mekalanos, J. J. (1987) Use of phoA gene fusions to identify a pilus colonization factor coordinately regulated with cholera toxin, Proceedings of the National Academy of Sciences. 84, 2833–2837.

