## Supplemental Figures S1 to S5 and Table S1 for "Peptide-based ligand antagonists block a *Vibrio cholerae* adhesin"

**Supplemental Information**

```
CLUSTAL O(1.2.4) multiple sequence alignment


FrhA_Split_PBD          SLDVKIVDDASTALDDRGEVDIVADSFTVSGVVANWTSWSNGTNVTTFDGTNAPNGGGLD       60
MpIBP_Split_PBD         --------SAPVAVDDQDSIVVQHEEFEVSEIAASWVSYTHGESVTTFDGTSD--LGGVD       50
                                .* .*:**:..: :  :.* ** :.*.*.*:::* .*******.    **:*

FrhA_Split_PBD          NDSGKDQIRWGQPASSYSSGYGFIDNDSALNGEFALNQDIILGTFTHYNYPVYSGGAITS       120
MpIBP_Split_PBD         NDSAKDQIRWGNPAESKQSGYGFIDNDSNLEGRFDLNQDISVGTFTHYNYPVYSGGAITS       110
                        ***.*******:**.* .********** *:*.* ***** :******************

FrhA_Split_PBD          ASMDVAFSVTDAHGVLTPVTLKLNFDHNETPNTNNPEASKDIIKVGNTNVTFENAGALYT       180
MpIBP_Split_PBD         AEMSVEFSVLDHLGVSTPVTLTVNFDHNETPNTNDVNASRDIVTVQNTHVTFERDGDIYT       170
                        *.*.* *** *  ** *****.:***********: :**:**:.* **:****. * :**

FrhA_Split_PBD          LQVIGFRIPGT--NQIVTEIRTGENATNSYELVVRVGPGEGYELPSTSGNVLSNDVS---       235
MpIBP_Split_PBD         VQIVGFREVGNPDGEVVTSIYTNENAATSYELVVRVVEGDGYSLPSTEGNIFDDNGLGAD       230
                        :*::***  *.  .::**.* *.***:.********  *:**.****.**::.::

FrhA_Split_PBD          --GADVDMTVVGAASGNHVSSGVSGSVGSMIAGLYGNLILLADGSYTYQVTANASSIPND       293
MpIBP_Split_PBD         SLGADGSVTVVGVAVGAIVSS--NESVGHSIEGQYGNLVLNSDGSYVYDVTASVSDIPAG       288
                          *** .:****.* *  ***  . ***  * * ****:* :****.*:***..*.** .

FrhA_Split_PBD          AIEIFTYTMKDGDGDTSTALLSINVNRVTMADFNANQ      330
MpIBP_Split_PBD         ATESFAYLIQDQDGSTSSANLSINVGTN---------      316
                        * * *:* ::* **.**:* *****.
```

**Figure S1**: Protein sequence alignment of the PBD and split domains of FrhA with their homologs in *Mp*IBP. The sequences forming the domains were determined by X-ray crystal structures. Residues lining the ligand-binding pocket are indicated by red boxes. The sequences share 60.1% identity, and the residues lining the ligand-binding pocket are conserved. The alignment was generated by Clustal Omega on the EMBL-EBI Job Dispatcher [32, 33].

**GenicBio Limited**

**MASS SPECTROMETRY REPORT**

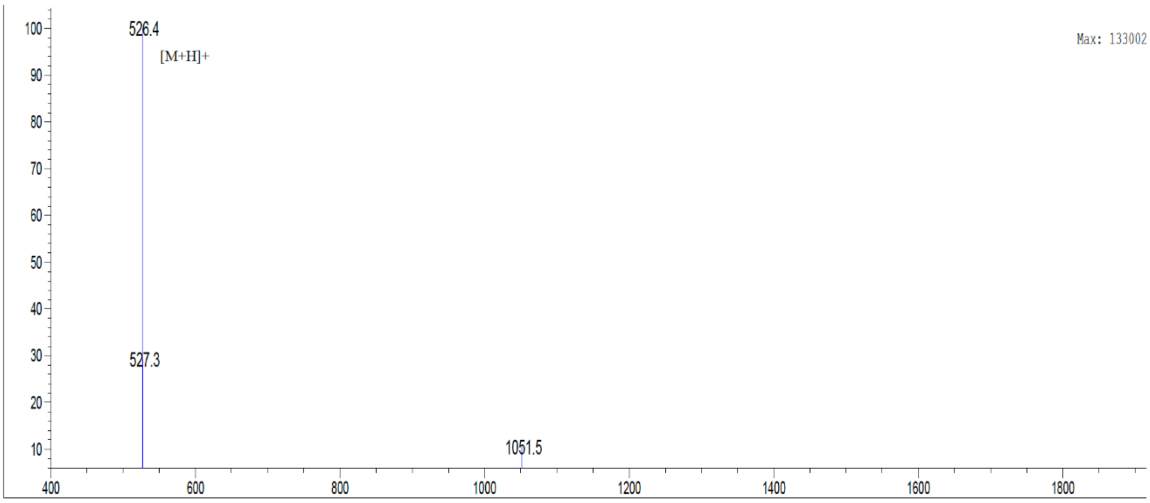


| | | | | | |
|---|---|---|---|---|---|
| Sample Description | | Instrument | Agilent-6125B | | |
| Analyzed date: | 2025-01-17 | Probe: | ESI | Probe Bias: | +4.5kv |
| Analyst: | Xiong | Nebulizer Gas Flow: | 1.5L/min | Detector: | 1.5kv |
| Sample: | Peptide 1 | CDL: | -20.0v | T. Flow: | 0.2ml/min |
| M.W.: | 525.57 | CDL Temp.: | 250°C | B. Conc.: | 50%H2O/50%ACN |
| Lot. No.: | P250113-GB728059 | Block Temp.: | 200°C | | |

**Figure S2**: Mass spectrum of the purified peptide AGYTD (GenicBio), confirming expected m/z peak. This report is representative of all peptides shown in this study. Mass spectra of all peptides are available upon request.

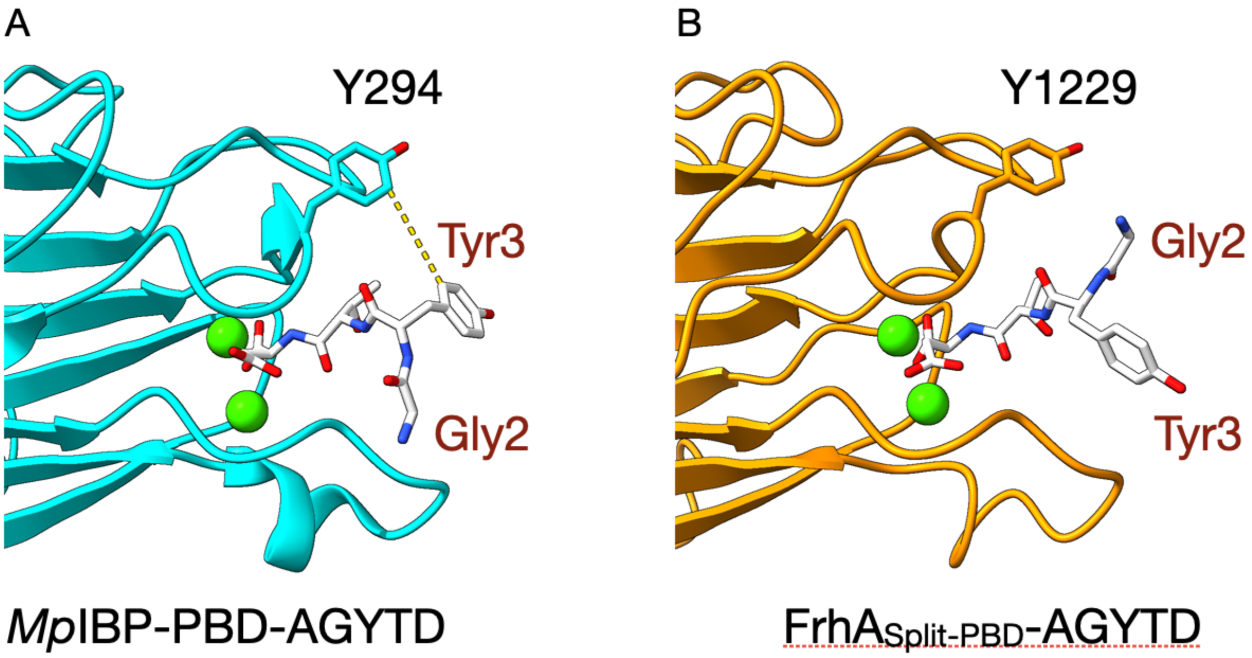

**Figure S3. Binding pockets of X-ray crystal structures of *Mp*IBP-PBD and FrhA$_{\text{Split-PBD}}$ binding to AGYTD.**

**(A–B)** Zoomed-in views of the ribbon models of the ligand-binding sites of AGYTD on *Mp*IBP-PBD and FrhA$_{\text{Split-PBD}}$. The peptide AGYTD is shown in white, with oxygen atoms in red and nitrogen atoms in blue. The $Ca^{2+}$ ions are depicted as green spheres. **(A)** The 1.6-Å X-ray crystal structure of *Mp*IBP-PBD (PDB ID: 6X5W) is shown in cyan [30]. The distance between the aryl side chains of tyrosine residues Tyr3 (peptide) and Y294 (*Mp*IBP-PBD) is shown as a yellow dashed line (~ 5 Å). **(B)** The X-ray crystal structure of FrhA$_{\text{Split-PBD}}$, the same as Fig. 1C, is shown in orange.

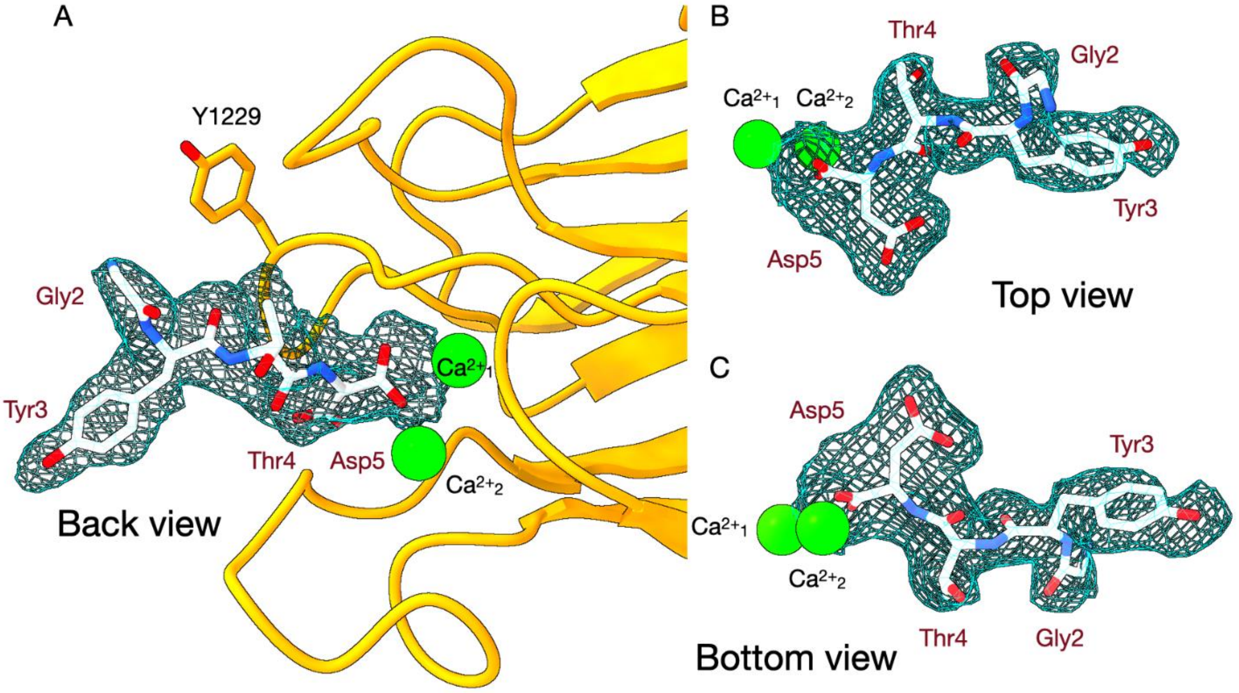


**Figure S4. Alternative views of the electron density of AGYTS in complex with FrhA$_{\text{Split-PBD}}$.**

**(A-C)** Views of the AGYTD peptide fitted in electron density, as in Fig. 1C (front view). The $Ca^{2+}$ ions are depicted as green spheres. The peptide AGYTS is shown in white, with oxygen atoms in red and nitrogen atoms in blue. **(A)** Back view, with FrhA$_{\text{Split-PBD}}$ shown in orange. **(B)** Top view. **(C)** Bottom view.

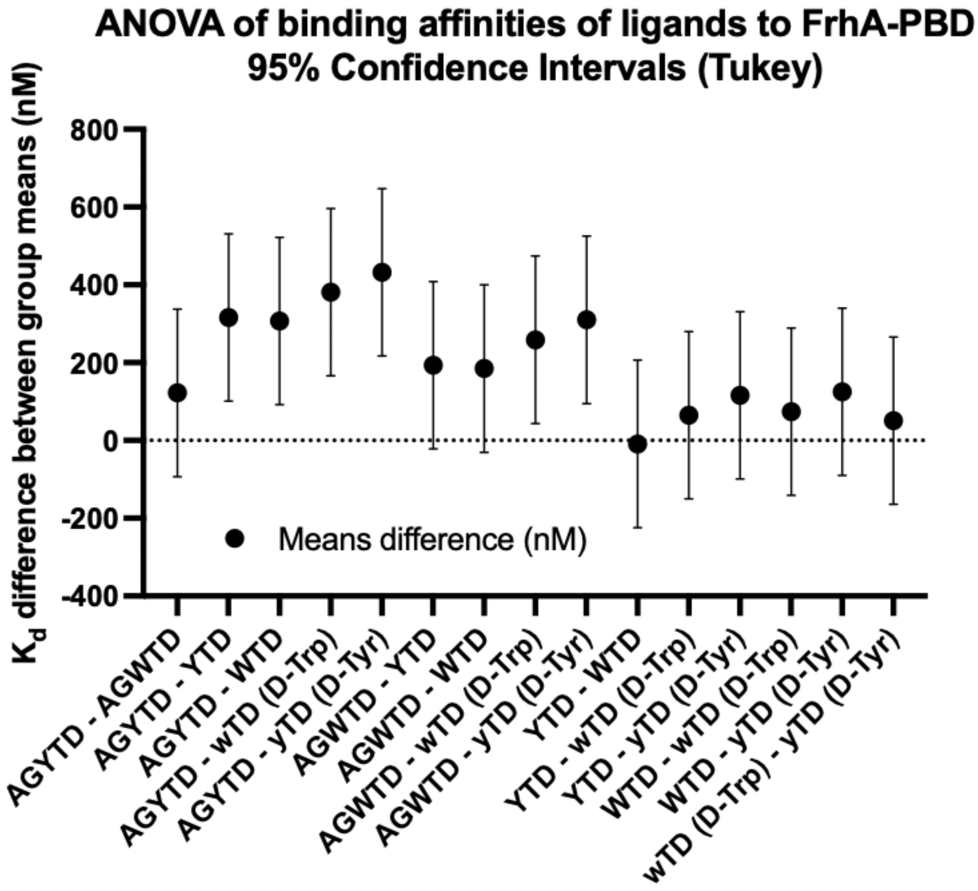


**Figure S5**: Plot of 95% confidence intervals of differences between binding affinities of ligands to FrhA-PBD according to ANOVA tests. Significant difference is indicated by the error bars not in contact with the horizontal dotted line.

**Table S1: X-ray and refinement statistics***

| | $FrhA_{Split-PBD}$-AGYTD | $FrhA_{Split-PBD}$-AGWTD |
|---|---|---|
| **Data collection** | | |
| Wavelength | 0.97949 | 0.98400 |
| Resolution range [Å] | 59.18 - 2.4 (2.49 - 2.4) | 101.1 - 2.4 (2.49 - 2.4) |
| Space group | P 1 | P 1 |
| Unit cell dimensions [Å] | 61.394 118.717 118.728 | 61.273 118.726 118.706 |
| Unit cell angles [°] | 60.05 76.45 81.42 | 60.03 85.62 76.7 |
| Total reflections | 471360 (22047) | 980678 (45166) |
| Unique reflections | 140116 (6838) | 140275 (6929) |
| Multiplicity | 3.4 (3.2) | 7.0 (6.5) |
| Completeness (%) | 98.20 (97.76) | 98.63 (97.91) |
| Mean I/sigma(I) | 3.4 (0.8) | 5.6 (0.6) |
| Wilson B-factor | 21.99 | 31.44 |
| R-merge | 0.212 (0.8) | 0.196 (1.061) |
| R-meas | 0.252 (0.718) | 0.212 (1.154) |
| R-pim | 0.135 (0.396) | 0.080 (0.447) |
| CC1/2 | 0.904 (0.159) | 0.991 (0.700) |
| **Refinement** | | |
| Reflections used in refinement | 108400 (10808) | 108729 (10840) |
| Reflections used for R-free | 1549 (163) | 1567 (170) |
| R-work | 0.2531 (0.3328) | 0.2270 (0.2954) |
| R-free | 0.2589 (0.3678) | 0.2318 (0.2935) |
| Number of non-hydrogen atoms | 15774 | 15471 |
| macromolecules | 14001 | 14051 |
| ligands | 43 | 70 |
| solvent | 1730 | 1350 |

| | | |
|---|---|---|
| Protein residues | 1902 | 1908 |
| Nucleic acid bases | 0 | 0 |
| RMS (bonds) [Å] | 0.013 | 0.012 |
| RMS (angles) [º] | 1.38 | 1.28 |
| Ramachandran favored (%) | 95.44 | 97.76 |
| Ramachandran allowed (%) | 4.24 | 1.92 |
| Ramachandran outliers (%) | 0.32 | 0.32 |
| Rotamer outliers (%) | 5.27 | 4.16 |
| Clashscore | 5.46 | 2.56 |
| Average B-factor | 37.18 | 36.14 |
| macromolecules | 37.85 | 35.85 |
| ligands | 31.67 | 40.09 |
| solvent | 31.89 | 38.92 |

* Statistics for the highest-resolution shells are shown in parentheses.
